# Low MSH2 protein levels identify muscle-invasive bladder cancer resistant to cisplatin

**DOI:** 10.1101/359554

**Authors:** Andrew Goodspeed, Annie Jean, James C. Costello

## Abstract

**Background:** The response to first-line, platinum-based treatment of muscle-invasive bladder cancer has not improved in three decades.

**Objective:** The objective of this study is to identify genes that predict cisplatin resistance in bladder cancer.

**Design:** We performed a whole-genome, CRISPR-based screen in a bladder cancer cell line treated with cisplatin to identify genes that mediate response to cisplatin. Targeted validation was performed *in vitro* across two bladder cancer cell lines. The top gene candidate was validated in a publicly available bladder cancer dataset containing 340 bladder cancer patients with treatment, protein, and survival information.

**Results and limitations:** The cisplatin resistance screen suggested the mismatch repair pathway through the loss of MSH2 and MLH1 contribute to cisplatin resistance. Bladder cancer cells depleted of MSH2 are resistant to cisplatin *in vitro*, in part due to a reduction in apoptosis. These cells maintain sensitivity to the cisplatin-analog, oxaliplatin. Bladder tumors with low protein levels of MSH2 have poorer overall survival when treated with cisplatin- or carboplatin-based therapy.

**Conclusions:** We generated *in vitro* and clinical support that bladder cancer cell lines and tumors with low levels of MSH2 are more resistant to cisplatin-based therapy. Further studies are warranted to determine the ability of MSH2 protein levels to serve as a prospective biomarker of chemotherapy response in bladder cancer.

**Patient summary:** We report the first evidence that the protein level of MSH2 may contribute to chemotherapy resistance observed in bladder cancer. MSH2 levels has the potential to serve as a biomarker of treatment response.

## Introduction

Bladder cancer is the ninth most common cancer type with an estimated 430,000 new cases and 165,000 deaths worldwide in 2012 [1]. In addition to radical cystectomy, patients with muscle-invasive bladder cancer (MIBC) are typically treated with one of two chemotherapy regimens, MVAC (methotrexate, vinblastine, doxorubicin, cisplatin) or GC (gemcitabine, cisplatin), both of which contain cisplatin [2]. The response rate to these cisplatin-based chemotherapies in MIBC remains 50%, a rate that has seen little improvement over the last three decades [2]. Identifying the molecular mechanisms that determine patient response to these treatments has direct clinical impact, including as a biomarker to stratify patients according to response and potentially an avenue to explore novel therapeutic targets.

Several biomarkers of response to cisplatin-based chemotherapy have been proposed in bladder cancer. The most well-established and clinically validated biomarker is mutant ERCC2, which leads to defects in nuclear excision repair [3,4]. ERCC2 mutations are found in 9% of bladder cancer patients and suggest these patients will be exceptional responders to chemotherapy [5]. An alternative biomarker that identifies patients that are resistant to chemotherapy, rather than hypersensitivity, would supplement the use of ERCC2 mutations in the clinic. This type of biomarker could alter standard-of-care by robustly identifying patients that will not respond to cisplatin-based chemotherapy and preemptively steer those patients to an alternative therapy.

Mediators of resistance to cisplatin have been described in other cancer types; for example, cancers deficient in the mismatch repair pathway are more resistant [6,7]. This occurs through a poorly understood mechanism that may include a combination of the DNA damage response [6], base excision repair [8], or translesion synthesis [9]. Several components of the mismatch repair pathway are altered in many cancer types, with the most frequently altered being MLH1 and MSH2 [10]. A subset of bladder cancers have reduced or absent protein expression of MSH2 [11–14]. These studies have reported inconsistent findings regarding the potential relationship between reduced MSH2 expression and various clinicopathologic features of bladder cancers. Additionally, none of these studies address how MSH2 may influence the response to chemotherapy despite mounting evidence from *in vitro* and acquired resistance studies performed in other cancer types [6,7].

Here, we take an unbiased approach to investigate mediators of cisplatin resistance by performing, to the best of our knowledge, the first genome-wide, cisplatin resistance screen using a CRISPR-Cas9 library in bladder cancer cells. Our screen suggests that cells with an MSH2 knockout are more resistant to cisplatin. This result was validated by showing bladder cancer cell lines with a MSH2 knockdown have a reduction in apoptosis following cisplatin treatment. Consistent with our *in vitro* work, we found tumors with low levels of MSH2 protein expression had a poorer response to cisplatin-based chemotherapy compared to patients with higher expression of MSH2.

## Materials and Methods

### Cell culture, shRNA knockdown, and drug treatments

MGHU4 and 253J bladder cancer cell lines were cultured in Minimum Essential Medium media (Gibco) supplemented with 10% fetal bovine serum (Sigma-Aldrich). PLKO.1 TRC shRNAs were transduced using psPAX2 (Addgene plasmid # 12260) and pMD2.G (Addgene plasmid # 12259) (Supplementary Table 1) [15]. Transduced cells were selected with 2-5 µg/mL puromycin (Sigma-Aldrich). Cisplatin (Sigma-Aldrich), methotrexate (ApexBio), adriamycin (ApexBio), vinblastine (ApexBio), and oxaliplatin (ApexBio) were solubilized in DMSO, and gemcitabine (Sigma-Aldrich) in water. For dose response experiments, cell viability was measured using the CellTiter-Glo™ luminescent assay (Promega).

### Performing the CRISPR resistance screen

To generate sgRNA lentivirus, 12µg of the human GeCKO (Genome-Scale CRISPR Knock-Out) lentiviral A library was transfected into 30 million HEK293T cells for 24 hours [16,17]. Viral supernatant was harvested and MGHU4 cells were transduced at a calculated multiplicity of infection of 0.3. Cells were selected with 2-5 µg/mL puromycin. Two million MGHU4 cells were plated in quadruplicate for each condition. Cells were treated with DMSO or 3mM cisplatin for 30 hours. Treatment media was removed and cells were allowed to grow to confluency prior to harvesting genomic DNA (Machery-Nagel).

### Sequencing of sgRNA

The sgRNA sequences of each replicate were PCR amplified from 4µg of genomic DNA using primers containing adaptor and barcoding sequences. The resulting DNA fragments were separated using agarose gel electrophoresis and purified using gel extraction (Machery-Nagel). DNA fragments were sequenced using a 1X125bp run on the HiSeq 2500 (Illumina). Individual sgRNA sequences were indexed using the bowtie2-build function. Reads generated from each sample were aligned to the indexed sgRNA sequences with the bowtie2 sequence aligner using the “very-sensitive-local” option [18]. sgRNA counts for each sample were summarized using htseq [19]. sgRNA counts across all samples were compiled and differential sgRNA abundance was calculated using DESeq2 [20]. A heatmap of raw reads was generated using gplots [21]. To map the sgRNAs results to the gene level (~3 gRNAs per gene), we calculated the mean fold change and combined p-values using Fisher’s method followed by multiple hypothesis testing correction using the Benjamini-Hochberg procedure [22].

### Resistance screen enrichment analysis

The Database for Annotation, Visualization and Integrated Discovery (DAVID) was used to determine enriched GO term biological processes [23,24]. The top resistant genes were identified using the thresholds of a combined q-value <0.01 and log_2_ fold change >2 in the cisplatin-treated group compared to the DMSO control group, resulting in 48 resistant genes. In addition to GO term enrichment, Gene set enrichment analysis (GSEA) was used to determine the enrichment of the KEGG Mismatch Repair pathway across the entire gene list ranked by mean fold change of the sgRNAs for each gene [25–27].

### Western blots

A total of 25µg of protein was loaded onto a 10% SDS-PAGE and transferred to nitrocellulose membranes (BioRad). Membranes were blocked with 5% bovine serum albumin in Tris-buffered saline with 0.1% tween 20 (TBS-T) (Sigma-Aldrich). Membranes were incubated overnight at 4°C (see Supplementary Table 2 for antibodies). Following TBS-T washes, membranes were incubated with secondary antibody at room temperature for 1 hour. Membranes were imaged using the Odyssey Fc Imaging System (LI-COR) following application of ECL (Millipore Sigma).

### Caspase activation

Bladder cancer cell lines were co-treated with the indicated concentration of cisplatin and 4µM CellEvent™ Caspase-3/7 Green Detection Reagent (Thermo Fisher Scientific) according to manufacturer instructions. Phase contrast and GFP were imaged every 3 hours following initial treatment using the Incucyte Zoom system (Essen Bioscience) at a 4X objective. The displayed results are cumulative GFP counts normalized to total cell confluency.

### qRT-PCR

Total RNA was extracted using the NucleoSpin RNA kit (Machery-Nagel). cDNA synthesis was performed with the iScript cDNA Synthesis Kit (BioRad). The SsoFast Evagreen supermix (BioRad) was used for qRT-PCR assays (see Supplementary Table 3 for primer sequences).

### Analysis of bladder tumors

All MIBC data was generated by The Cancer Genome Atlas (TCGA) [28]. Clinical data containing tumor characteristics, patient treatments, and survival was downloaded using the TCGAbiolinks R package [29]. Level 4, batch normalized Reverse phase protein array (RPPA) for all TCGA bladder cancer samples was downloaded from The Cancer Proteome Atlas [30]. MSH2 mRNA (RNA Seq V2 RSEM) expression was downloaded from cBioPortal for Cancer Genomics [31,32]. TCGA bladder cancer patients receiving cystectomies were identified using data from the Broad GDAC Firehose [33]. Finally, we also used published neoantigens for TCGA bladder cancer samples [5]. Overall, 340 TCGA patients had RPPA and some form of clinical data available (Supplementary Table 4). Overall survival was analyzed and plotted using the survival, survminer, and ggplot2 R packages [34–36]. Statistical significance between two groups in survival analysis was performed using a log-rank test.

## Results

### Whole-genome CRISPR screen identifies mediators of cisplatin resistance in a bladder cancer cell line

We executed an unbiased CRISPR screen to identify mediators of cisplatin resistance in a bladder cancer cell line. In contrast to synthetic lethal relationships, we identified the sgRNA constructs that gain in relative number in the cisplatin compared to DMSO treated groups. MGHU4 cells were transduced with a genome-wide CRISPR library consisting of 65,386 sgRNAs (~3 sgRNAs per gene) (Figure 1A) [16,17]. Transduced MGHU4 cells were treated with either DMSO or cisplatin for 30 hours. The sgRNA constructs that change in abundance in the cisplatin compared to the DMSO treated group indicate an impact on cisplatin sensitivity (Supplementary Table 5). Using median-centered raw reads, we identified 9,757 sgRNA constructs that were differentially abundant between the two groups (least fold-change ≥ 3) (Figure 1B). These constructs were further filtered to the 56 sgRNA constructs that increased in relative abundance in the cisplatin compared the DMSO control group (Figure 1C and 1D). We found that MSH2 and MLH1 were the top two gene candidates based on statistical significance (Figure 1E, Supplementary Table 6).

**Figure 1.**
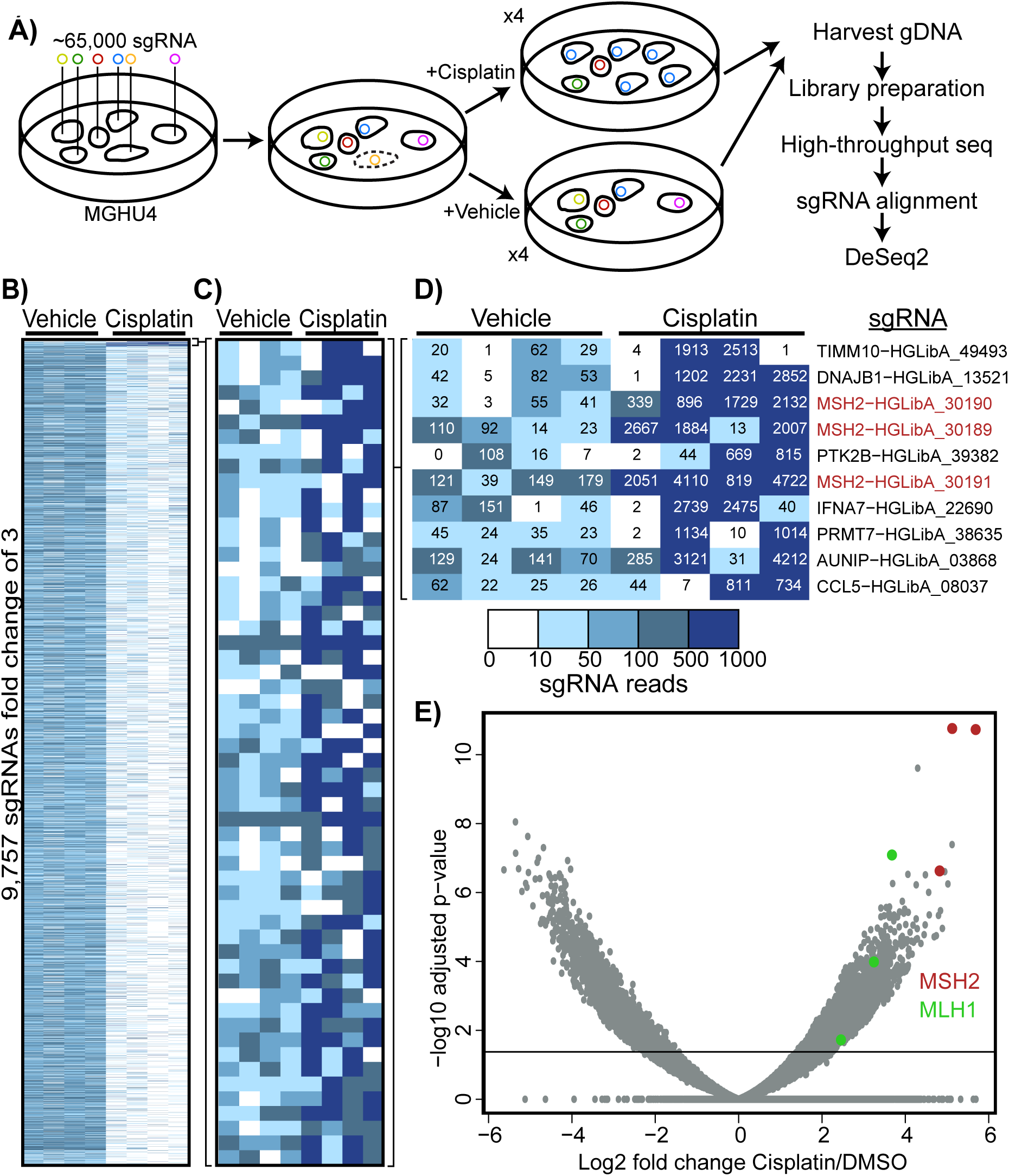
A whole-genome CRISPR screen to identify mediators of cisplatin resistance in a bladder cancer cell line. (A) Experimental outline of the screen and analysis. (B) A heatmap displaying median-centered counts for 9,757 differentially abundant sgRNAs. (C) The median-centered counts of the 56 sgRNAs more abundant in the cisplatin-treated group. (D) Counts from the screen for the top 10 resistant sgRNAs. (E) A volcano plot displaying the fold-change and adjusted p-value for all sgRNAs identified in the screen. A threshold of 0.05 adjusted p-value is indicated on the plot.

### The mismatch repair pathway is a mediator of cisplatin sensitivity

We performed an unbiased pathway-level analysis to identify processes that mediate cisplatin resistance. Gene Ontology (GO) term enrichment analysis was performed on the top 48 resistant genes from our screen [23,24]; the top pathway was mismatch repair (Figure 2A). To confirm this result, we tested the mismatch repair pathway using GSEA on the mean fold change of all 16,564 genes identified in our screen and found it was significantly enriched in cisplatin resistance (p < 0.05) (Figure 2B). MLH1 and MSH2 each had 3 significantly resistant sgRNA constructs and several other genes in the mismatch repair pathway had a single resistant sgRNA (Figure 2C).

**Figure 2.**
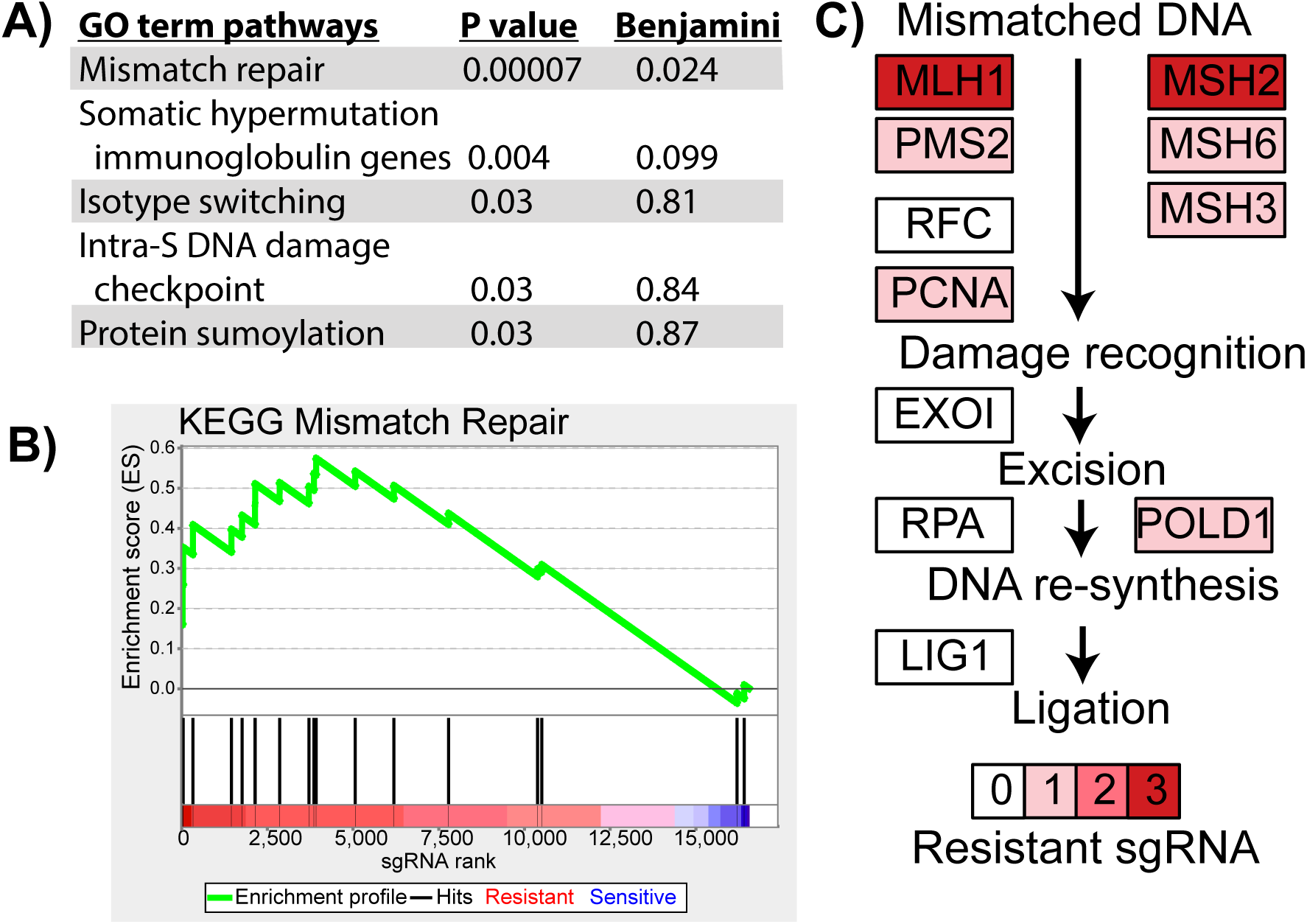
sgRNA constructs targeting genes of the mismatch repair pathway are enriched in the cisplatin-treated MGHU4 cells. The mean fold change and combined p-value was calculated using the 3 sgRNAs targeting each gene in the screen. (A) The top GO term biological processes results from DAVID enrichment analysis of the 48 genes in the screen with a combined q-value <0.01 and log _2_mean fold change >2 of the cisplatin versus DMSO group. (B) The GSEA results of the KEGG mismatch repair pathway on the ranked list of genes by mean fold change. (C) The components and genes of the mismatch repair pathway are depicted with the number of significantly resistant sgRNAs from our synthetic lethal screen targeting each gene (out of 3).

### Bladder cancer cell lines with knockdown of MSH2 are resistant to cisplatin

Our screen suggests that the loss of mismatch repair offers resistance to cisplatin in bladder cancer cell lines, consistent with findings in other cancer types [6,7]. To confirm our finding, we found that strong shRNA knockdowns of MSH2 provided roughly two-fold resistance to cisplatin in MGHU4 and 253J bladder cancer cell lines (Figure 3A-3C). We also found that these cells had reduced caspase activation compared to non-targeting control cells when treated with cisplatin (Figure 3D-3G). Collectively, these data show that the loss of MSH2 in cisplatin-treated bladder cancer cells increases survival and reduces apoptosis.

**Figure 3.**
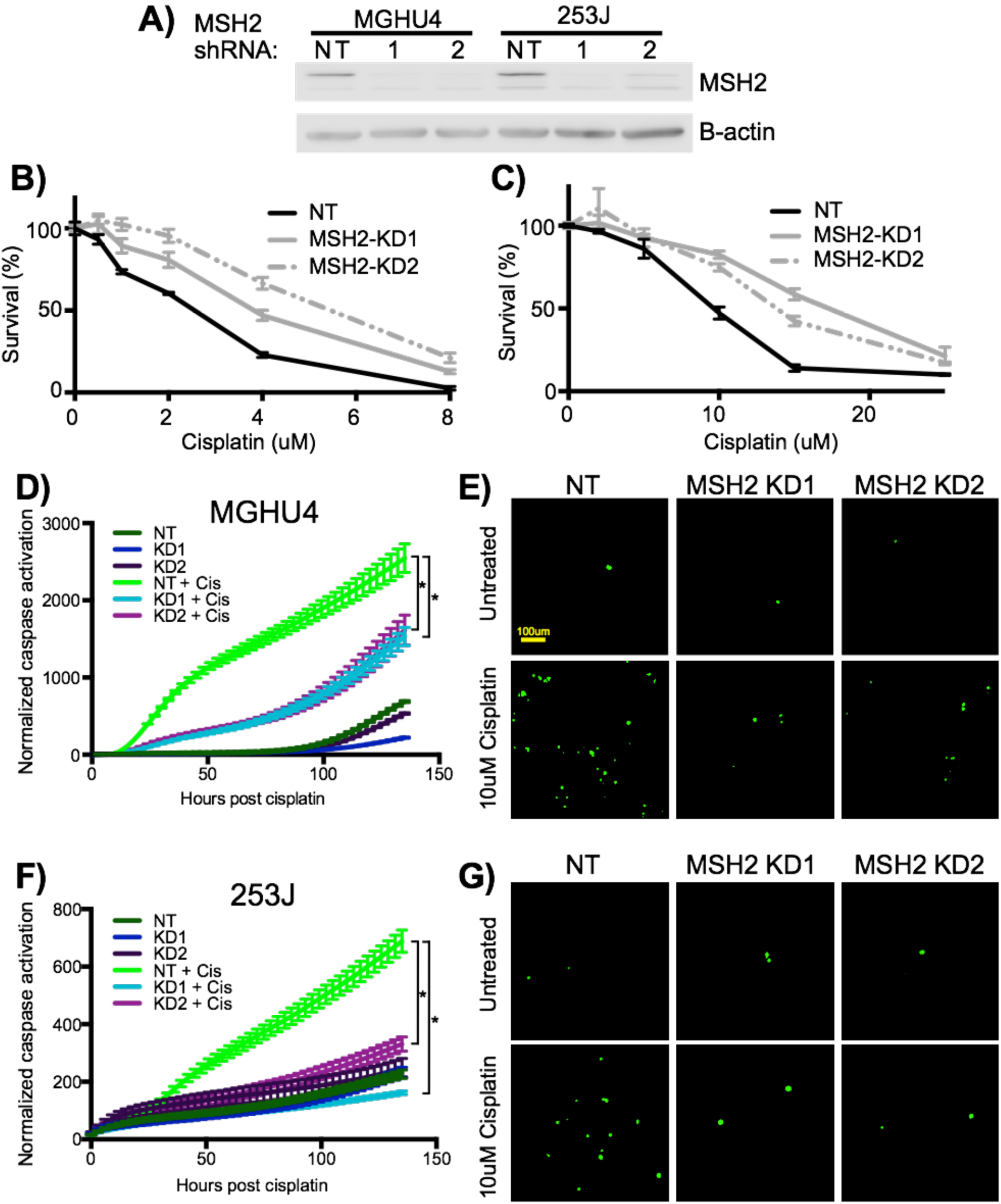
Knockdown of MSH2 increases cisplatin resistance in bladder cancer cell lines. Two shRNA constructs targeting MSH2 and one non-targeting shRNA control were transduced into MGHU4 and 253J bladder cancer cell lines. (A) Western blot showing the level of knockdown of MSH2. 253J (B) and MGHU4 (C) cell lines were treated with the indicated doses of cisplatin for 48 hours and cell viability was measured using an ATP-based assay. MGHU4 (D) and 253J (F) cells were treated with 10µM and 15µM cisplatin respectively or vehicle. Cumulative caspase activation (GFP+ cells) was measured and normalized to cell confluency. Representative images of MGHU4 (E) and 253J (G) cells are shown at the 24 hour time point. A two-way ANOVA was performed on the mean ± SEM of a representative experiment (of 3 experiments).

### MSH2 knockdown impairs cisplatin-induced DNA-damage response

Several studies have indicated that a lack of DNA-damage response is responsible for cisplatin resistance in mismatch repair deficient cells. p73 [6] and p53 [37] have been shown to be stabilized in a mismatch repair-dependent manner following cisplatin. However other work has shown that p53 is activated independent of mismatch repair downstream of cisplatin [6]. Additionally, phosphorylation of ATM has been shown to be upstream of mismatch repair and necessary for the activation of p73 and p53 [37,38].

To investigate the role of the DNA-damage response, we treated MGHU4 and 253J MSH2 knockdown cells with cisplatin for 0, 2, 24, and 48 hours followed by a western blot and qRT-PCR to look at factors relating to DNA damage (Figure 4). As expected, cisplatin treatment in MGHU4 and 253J cells transduced with non-targeting shRNA leads to an accumulation of p-ATM and p53, but these bladder cancer cell lines did not express measurable levels of p73. We hypothesized that cells depleted of MSH2 will have a significant attenuation of the DNA-damage response. We observed reductions in the accumulation p-ATM and stabilization of p53. Cisplatin also leads to transcriptional induction of BAX, MDM2, p21, and PUMA in all cells tested, but induction of BAX and MDM2 is slightly attenuated in cells with a MSH2 knockdown. In the 253J cells, induction of p21 is also attenuated. Collectively, we observe a stronger DNA-damage response in MSH2 knockdown cells than has been observed in past studies but there is still an impairment in some elements of the DNA-damage response [6,37,38].

**Figure 4.**
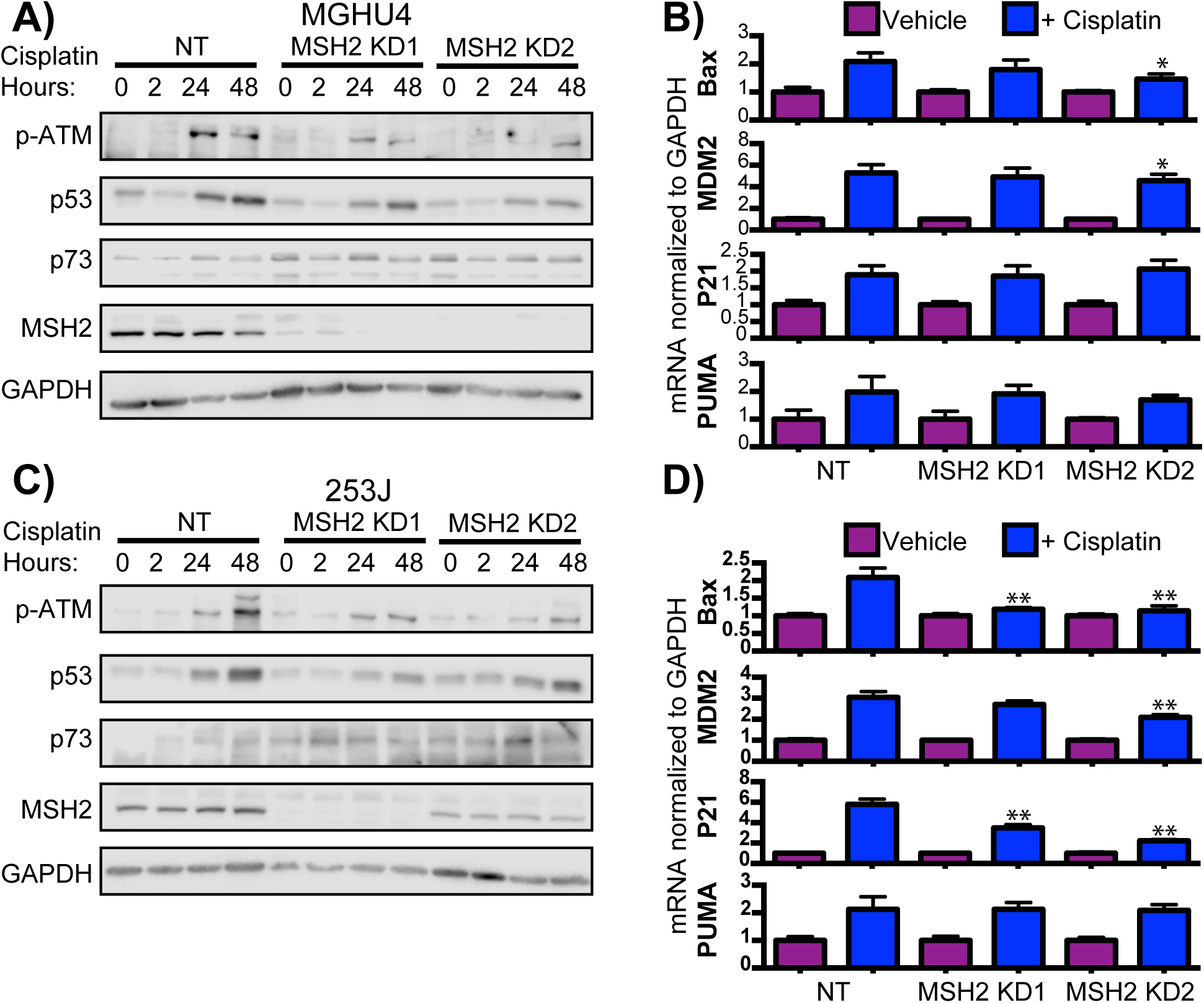
The DNA-damage response is slightly reduced in bladder cancer cell lines with knockdown of MSH2 compared to non-targeting controls treated with cisplatin. MGHU4 (A) and 253J (C) bladder cancer cell lines containing a non-targeting or one of two MSH2 shRNA constructs were treated with 2µM cisplatin for 0, 2, or 24 hours and a western blot was performed. MGHU4 (B) and 253J (D) bladder cancer cell lines were treated with 6µM and 12µM cisplatin for 24 hours and qRT-PCR was performed. The indicated genes were normalized to GAPDH expression and a two-way ANOVA was performed on the mean ± SEM of a representative experiment (of 3 experiments).

### Cells with low levels of MSH2 are equally sensitive to oxaliplatin and other chemotherapies

Previous work has shown that the loss of mismatch repair does not deter sensitivity to the cisplatin analog, oxaliplatin [39]. In bladder cancer, oxaliplatin has been shown to have similar activity to cisplatin [40–42]. Therefore, we tested whether oxaliplatin as well as the other constituent components of the MVAC and GC regimens are effective in bladder cancer cell lines with reduced MSH2 expression. We found that MGHU4 and 253J cells with reduced MSH2 were equally sensitive to all of the tested chemotherapies compared to control (Figure 5, Supplementary Figures 1 and 2). This indicates that MSH2 loss mediates resistance to cisplatin specifically, but MSH2 loss does not affect the other drugs in the standard MVAC and GC treatments or the cisplatin analog, oxaliplatin.

**Figure 5.**
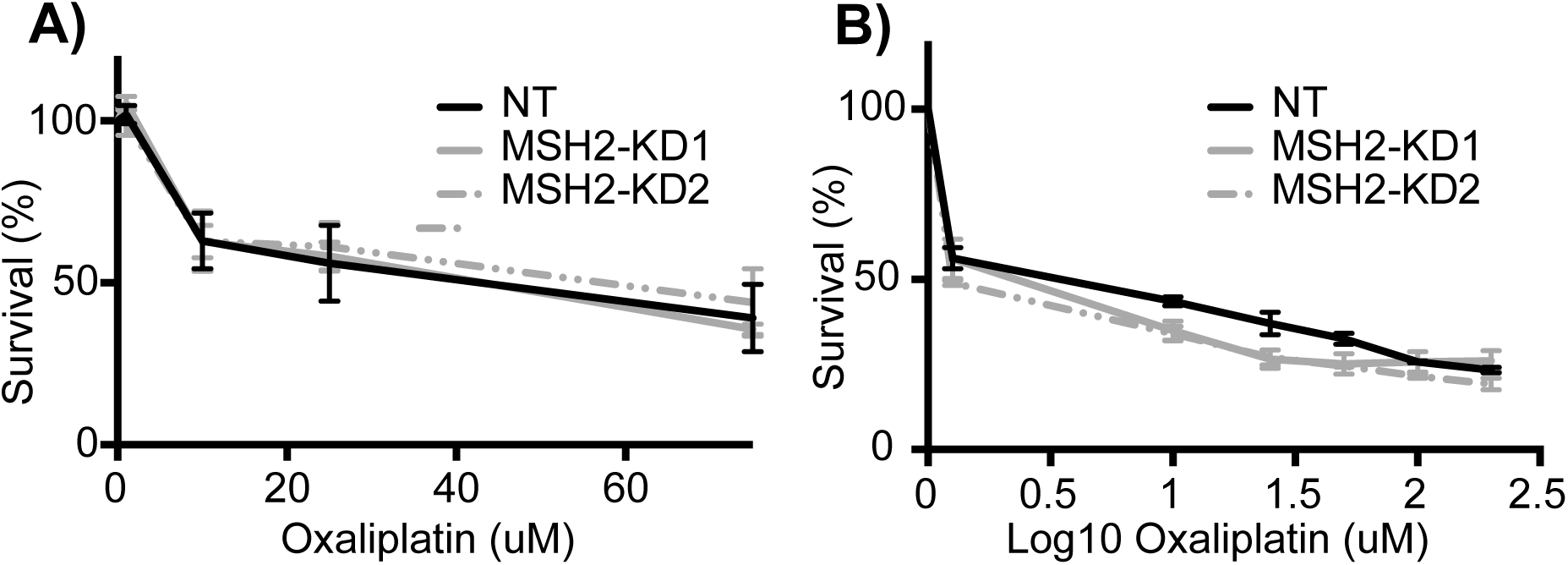
MSH2 knockdown bladder cancer cell lines are equally sensitive to oxaliplatin. MGHU4 (A) and 253J (B) bladder cancer cell lines were treated with the indicated doses of oxaliplatin for 48 hours and cell viability was measured using an ATP-based assay. No difference in response was observed between control and knockdown of MSH2.

### MSH2 protein levels do not correlate with clinicopathologic features in bladder cancer

To investigate the clinical impact of MSH2 in bladder cancer, we analyzed 340 bladder cancer patients from TCGA that had both molecular and clinical data [28]. We first hypothesized that MSH2 and MSH6 protein levels would strongly correlate because they form the MutSα mismatch repair complex and the presence of one influences the stability of the other [43]. We found that MSH2 and MSH6 protein levels, as measured by RPPA, did strongly correlate with each other (Supplementary Figure 3A, *ρ* = 0.66). We found only a very weak correlation between MSH2 mRNA and protein levels, indicating that mRNA levels are a poor surrogate for MSH2 protein levels in bladder cancer (Supplementary Figure 3B, *ρ* = 0.29). Deficiency in the mismatch repair pathway typically leads to an increase in mutations. Surprisingly, low MSH2 protein levels did not have an impact on the total number of neoantigens in bladder cancer (Supplementary Figure 3C) [5]. Additionally, MSH2 protein levels are unchanged with regards to tumor stage or grade, suggesting MSH2 is not prognostic of clinicopathologic features (Supplementary Figure 3D-3E).

### MSH2 levels correlate with the response to platinum-based chemotherapy in bladder cancer patients

We investigated if MSH2 protein levels correlate with the response to cisplatin- or carboplatin-based (platinum-based) chemotherapy in bladder cancer patients. First, we divided all TCGA bladder cancer patients into 3 groups based on their MSH2 protein expression (Supplementary Figure 4A). We found that MSH2 levels were significantly higher in bladder cancer patients who had a complete or partial response to platinum-based chemotherapy compared to those with stable or progressive disease (Supplementary Figure 4B and Figure 6A, p = 0.0003, Wilcoxon rank-sum test). The majority of patients with tumors having high or medium levels of MSH2 had a complete response to platinum-based therapy while the majority of patients with low MSH2 experience had progressive disease (Figure 6B).

**Figure 6.**
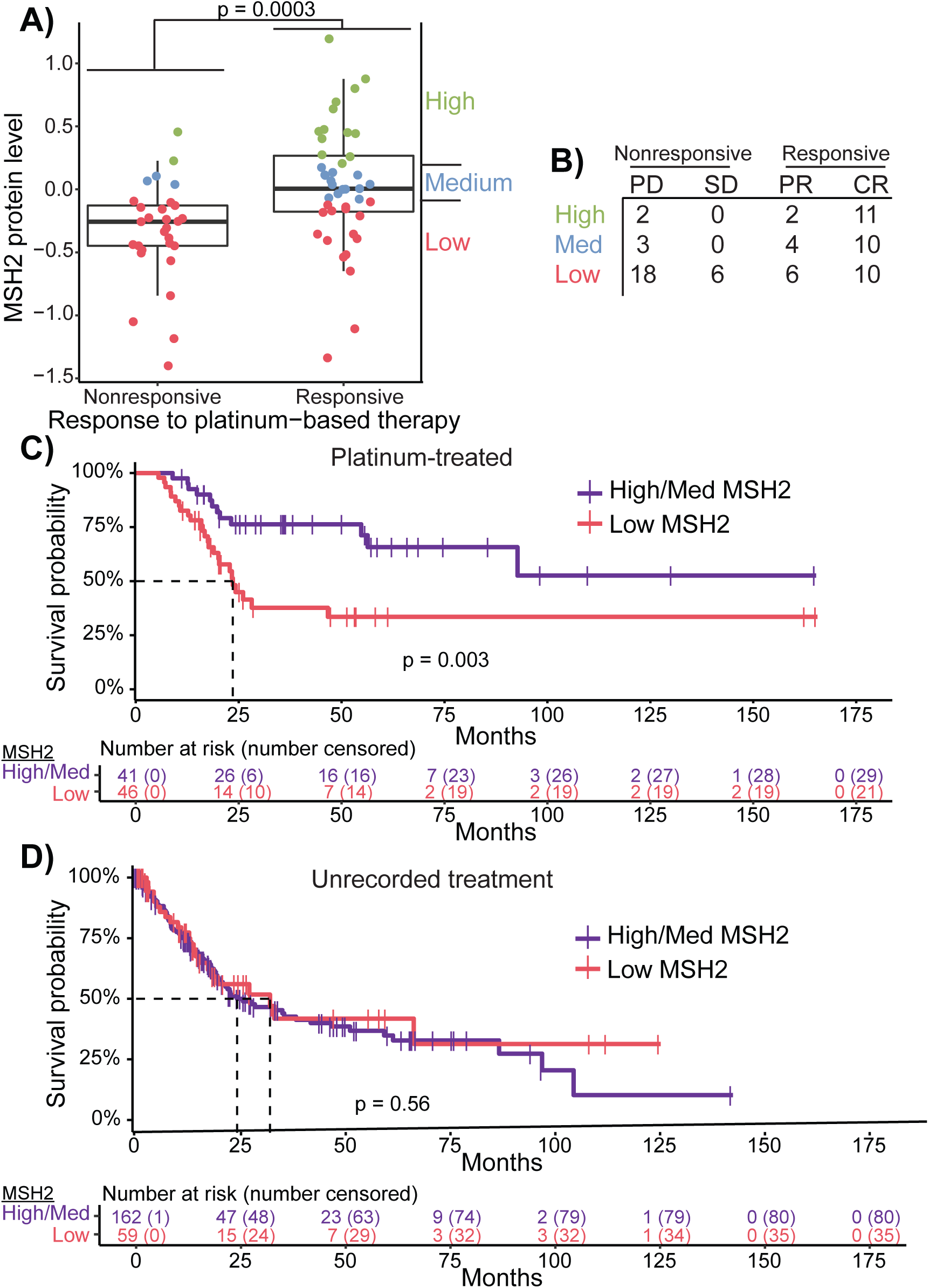
Bladder cancer patients with low levels of MSH2 protein correlate with a poorer response to platinum-based chemotherapy. The response to platinum-based chemotherapy was compared to the MSH2 RPPA levels in 340 bladder cancer patients. (A) MSH2 protein levels were plotted according to their response to platinum-based chemotherapy (PD, progressive disease; SD, stable disease, PR, partial response; CR, complete response). The statistical difference between good (complete (CR) and partial (PR) response) and poor (stable disease (SD) and progressive disease (PD)) responders was calculated using a Wilcoxon rank-sum test (P = 0.0003). Each point is colored by MSH2 protein level: low (red), medium (blue), or high (green). (B) Table showing the number of patients of each MSH2 group by type of response. Overall survival is plotted for bladder cancer patients with a platinum-based treatment (C) or a non-pharmacologic or radiation treatment (D). Survival of patients with low MSH2 (red) is compared to those with medium or high levels of MSH2 (purple). The statistical difference in survival was calculated using a log-rank test.

We next asked how MSH2 protein levels affect the overall survival of bladder cancer patients. We found that for patients treated with platinum-based therapy, the individuals with low MSH2 had significantly poorer overall survival compared to patients with medium or high protein levels (Supplementary Figure 4C and Figure 6C, p = 0.003, log-rank test). In contrast, when patients that did not have a recorded pharmacologic or radiation therapy are considered, those with low MSH2 have equivalent survival rates compared to patients with medium or high MSH2 levels (Supplementary Figure 4D and Figure 6D, p = 0.56, log-rank test). Importantly, the patients in different MSH2 groups had no discernable differences in age, tumor stage, lymph node infiltration, metastasis, or cystectomy procedures (Supplementary Tables 7-9).

Lymph node positive and higher stage bladder cancers are strong prognostic factors of poorer overall survival [44]. We asked whether MSH2 protein levels still serve as a biomarker in this high-risk population. Similar to the larger population, we found that patients with low MSH2 had poorer overall survival when compared to other platinum-treated patients, but not when compared to other patients without a recorded treatment (Supplementary Figure 5A and 5B). Collectively, these results suggest that clinical response to platinum-based chemotherapy and subsequent overall survival are poorer in bladder cancer patients with low MSH2 protein levels.

## Discussion

No clinically actionable biomarker of resistance to first-line therapy in MIBC is available [3,4]. The mismatch repair pathway has been shown to impact the response to cisplatin-based chemotherapy in other cancers [6,7]. Here we report the first association of mismatch repair predicting response to cisplatin-based treatments in bladder cancer.

To the best of our knowledge, we performed the first genome-wide screen to identify mediators of cisplatin treatment in a bladder cancer cell line. We identified MSH2 and validated that bladder cancer cell lines with knockdown of MSH2 are resistant to cisplatin-mediated apoptosis. Similar to studies in other cancer types [6,37,38], we only observe reductions in the DNA damage response when cells have low levels of MSH2. It is unclear that reduction in the DNA-damage response alone accounts for the cisplatin resistance we observe, or if other components also contribute. Future studies are needed to address this potential link, and also determine if the DNA-damage response can serve as a therapeutic target to address mismatch repair-mediated cisplatin resistance.

While microsatellite instability is infrequent in bladder cancer [12,13], a subset of bladder cancers have low to no expression of MSH2 as determined by immunohistochemistry [11–14]. Using RPPA data from the TCGA, we found that the one third of patients expressing low levels of MSH2 protein have an impaired response to platinum-based chemotherapy. This cohort has previously been analyzed for MSI and only a single patient was been found to be MSI-high [45]. Therefore, roughly one third of MIBC patients appear to have tumors with reduced MSH2 expression, that may impact the response to platinum-based therapy, but do not have MSI or an increase in neoantigens (Supplementary Figure 3C). Future work should investigate the potential separation of mismatch repair functions in regards to DNA repair activity versus the response to cisplatin. Even though bladder cancers are typically microsatellite stable, defects in mismatch repair protein expression may still impact the response to cisplatin-based therapy.

The response rates to oxaliplatin-based therapy in bladder cancer range from 36-48% [41,42] in patients unfit for cisplatin and a disease-control rate of 36% in patients who have failed cisplatin- or carboplatin-based chemotherapy [40]. These studies demonstrate the usefulness of oxaliplatin in treating MIBC. We found that oxaliplatin was equally effective in bladder cancer cell lines who have regular or reduced MSH2 expression, despite these cell lines being resistant to cisplatin (Figure 3 and Supplementary Figure 1). We also showed that the non-cisplatin components of the MVAC and GC chemotherapy regimens are equally effective in MSH2 knockdown bladder cancer cell lines. Therefore, an important remaining question is if bladder tumors with low MSH2 levels would respond well to MVAC and GC regimens with oxaliplatin being supplemented for cisplatin.

## Conclusions

In this study, we identified that MSH2 and the mismatch repair pathway predict response to cisplatin-based chemotherapy in bladder cancer. Oxaliplatin maintains efficacy in MSH2 low bladder cancer cell lines suggesting that the substitution of cisplatin for this agent may serve as an effective alternative therapy. Future studies will be necessary to validate MSH2 protein levels as a prospective biomarker of platinum-based therapeutic resistance in bladder cancer.

## Acknowledgements

We would like to thank the other members of the lab, particularly Robert T Jones and Rani Powers for their special insight. We would also like to thank the Functional Genomics Facility for genomic library expertise.

